# Widespread Regulatory Turnover Across Human Segmental Duplications

**DOI:** 10.64898/2026.01.21.700947

**Authors:** Alexis Morrissey, Alan Brown, Jianyu Yang, Shaun Mahony

## Abstract

Segmental duplications (SDs) are a pervasive feature of eukaryotic genomes, enabling genomic innovation via the duplication of genes and *cis*-regulatory elements. However, the regulatory mechanisms that govern these regions of the genome after duplication are not well understood. The repetitive nature of SDs pose problems for short-read assays such as ChIP-seq and DNase-seq as they create reads that map to more than one location along the genome. Moreover, the newest Telomere-to-Telomere (T2T) human genome assembly has revealed previously unknown segmental duplications. In this study, we used the Telomere-to-Telomere genome assembly along with the probabilistic allocation of multi-mapped reads by our software Allo to better understand the regulatory mechanisms governing segmental duplications in the human genome. Using 134 transcription factor ChIP-seq datasets in GM12878, we found that transcription factor binding sites within segmental duplications are rarely conserved across paralogous copies. Similarly, chromatin states defined via histone ChIP-seq datasets are also poorly conserved across segmentally duplicated regions. In contrast, paralogous genes within SDs exhibit highly correlated expression patterns. Hi-C analysis revealed that many SD paralogs have higher than expected spatial proximity, suggesting that three-dimensional genome organization may buffer transcriptional divergence following duplication despite the loss of *cis*-regulatory regions. Together, our results suggest that regulatory regions undergo rapid turnover following segmental duplication, but the regulatory impact may be buffered by spatial proximity within the nucleus. Tissue-specific genes and transcription factors showed greater divergence, highlighting the role of SDs in expanding specialized cellular functions.

## Introduction

Gene duplication is a common mechanism by which new genes are created and is thus an important driver of genome complexity and evolution^1,2^. There are numerous ways in which duplicated genes can arise. A significant portion of the genome can be duplicated simultaneously, as in the case of whole genome duplication or chromosomal duplications. Smaller portions of the genome may also be duplicated through segmental duplications (SDs) or transposable element activity. In each case, duplicated genes have proven to be advantageous. For example, there is evidence that a whole genome duplication (WGD) took place at least once in the evolution of *Saccharomyces cerevisiae*^3^ as well as evidence of two WGDs in the evolution of early invertebrates^4^. Polyploidy is also particularly pervasive in plant genomes and may play a role in their ability to rapidly evolve^5^. Smaller regional duplications, including segmental duplications, are also prevalent in genomes, constituting about 7% of the newest Telomere-to-Telomere (T2T) human genome assembly^6^.

There are multiple ways in which gene duplication can lead to adaptation and play a role in evolution. The most obvious mechanism is that gene duplication can allow for the increased dosage of a protein necessary in a new environment^7,8^. Additionally, duplicated genes that keep all or some of their ancestral functions are thought to shield the genome in the case of deleterious mutations in one or more copies. A prominent example of this was discovered in elephants whose cancer rates are much lower than expected due to the presence of 20 copies of the tumor suppressor TP53 within their genomes^9^.

It has been suggested that paralogs are more likely to share similar expression than have divergent expression^10^, as regulatory sequences proximal to the duplicated gene may also be duplicated. One study also found that paralogous genes have higher than expected rates of contact within the three dimensional genome^11^. These results suggest that there may be shared gene regulatory systems among paralogous gene families, including shared sets of transcription factor binding sites. However, some duplicated genes have been shown to have differing expression levels^12^. For example, duplicated gene expression can differ across tissue-types, suggesting a role in differentiation^13,14^. Aberrant expression of paralogous genes has also been shown to affect disease progression^15^.

Recently duplicated genic and non-coding regions pose a problem for regulatory genomics assays that rely on short read sequencing like ChIP-seq and DNase-seq, as they will produce multi-mapped reads that align equally well across conserved duplicated sequences. Most regulatory genomics pipelines discard multimapped reads, or assign them randomly to a valid mapping location. The resulting exclusion of duplicated regions in the human genome has resulted in a dearth of knowledge surrounding the gene regulatory networks of duplicated genes. This lack of information impacts recently duplicated genes the most as they have the highest sequence similarity. Segmental duplications are among the most recent duplications in the genome and have very high identity (>75%). In recent work, we introduced Allo, which accurately assigns multi-mapped reads to source regions using a unique combination of probabilistic read mapping and neural networks^16^. We also showed that segmentally duplicated regions were a significant source of multi-mapped reads in ChIP-seq datasets and that Allo can uncover previously uncharacterized regulatory signals in these regions. We hypothesized that many of the interactions between transcription factors and SDs have been largely ignored in previous analyses.

In this work, we use DNase-seq, ChIP-seq, RNA-seq, and Hi-C datasets from 23 cell types to investigate the conservation of gene regulatory networks within segmentally duplicated regions of the human genome. To enhance our analysis of duplicated regions, we used the new T2T human genome assembly and probabilistically allocated multi-mapped reads using Allo^16^. We show that while the sequence conservation is high in segmentally duplicated regions (>75%), the retention of transcription factor binding sites is low. We find a similar pattern when investigating histone modifications using chromatin state data. Finally, we find that spatial proximity of segmentally duplicated regions may buffer the loss of transcription factor binding and maintain the transcription of duplicated genes.

## Results

### The Telomere-to-Telomere human genome assembly and multi-mapped read allocation enable regulatory genomics analysis of segmental duplications

While transposable elements have been the most common focus when studying repeated regions, our previous findings suggested that duplicated genes are also a major source of multi-mapped reads in regulatory genomics assays^16^. Because DNase-seq and histone ChIP-seq cover the broadest range of genomic locations in our dataset, we used these assays to examine the prevalence of multi-mapped reads and to get an overall view of their genomic distribution. Similar to the findings in our previous study, we saw that multi-mapped reads made up a significant portion of the total reads in the DNase-seq and histone ChIP-seq datasets used in this study **(Figure 1A)**. We analyzed the source of multi-mapped reads in the aforementioned datasets and found that almost all multi-mapped reads originated from repetitive elements (REs) and segmentally duplicated regions (SDs) **(Figure 1B)**. With the inclusion of dual-annotated regions, up to 33% of multi-mapped reads may result from SDs. Fewer than 1% of the multi-mapped reads across all of these samples were not explained by either SDs or REs. We also found that the percentage of multi-mapped reads was dependent on the assay or histone modification being analyzed **(Figure 1A)**. H3K9me3 had significantly more multi-mapped reads than the other datasets we analyzed here. Generally, H3K9me3 is associated with heterochromatin and thus may overlap silenced SDs or REs^17^.

**Figure 1:**
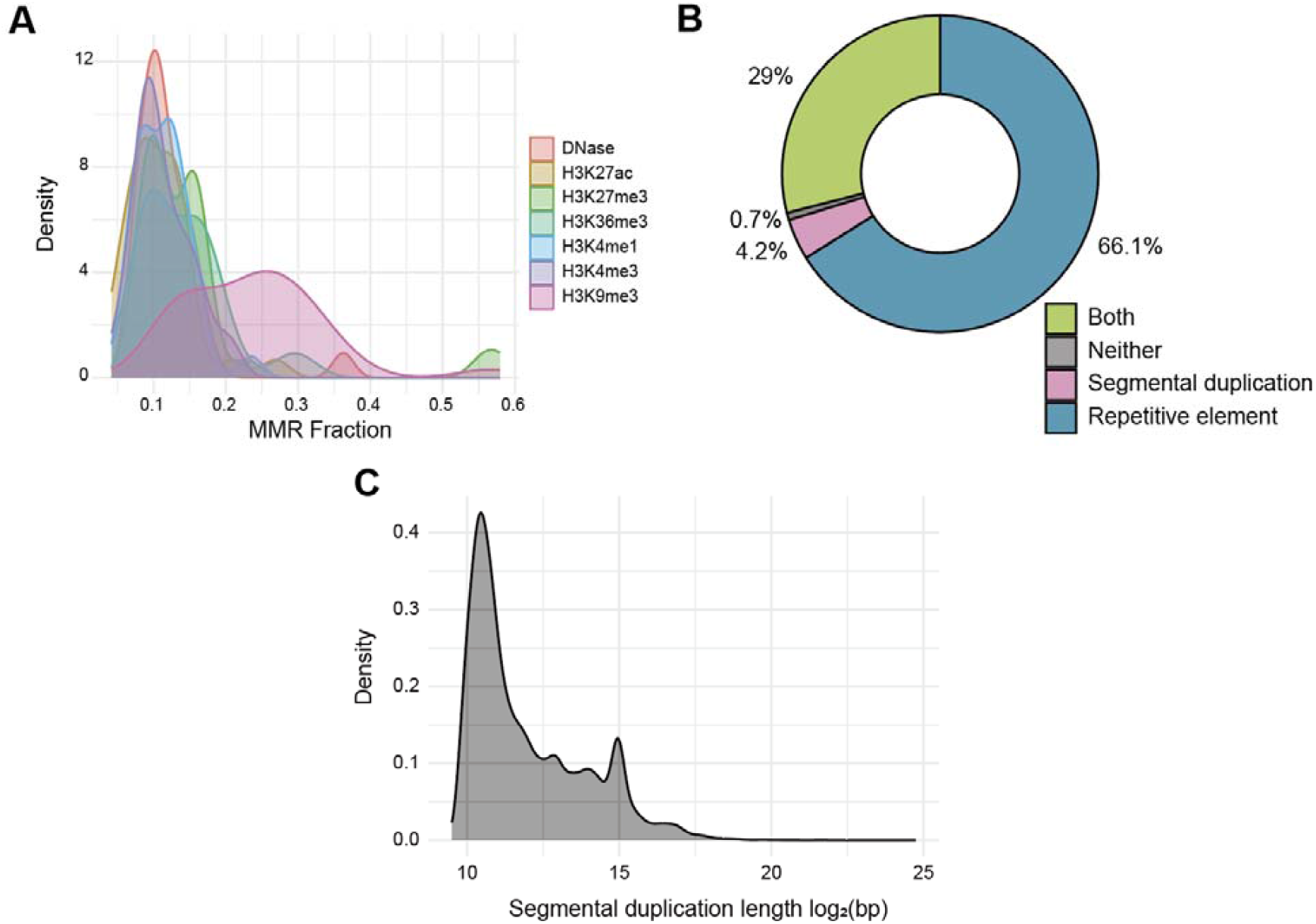
Proportions of multi-mapped reads in DNase-seq and ChIP-seq data. A) Density plot of the percentage of multi-mapped reads (MMRs) across DNase-seq and ChIP-seq histone datasets. B) Donut plot showing the percentage of multi-mapped reads that originated from repetitive elements, segmental duplications, or both. **C)** Length of segmental duplications as called by BISER in the T2T genome assembly.

The newest version of the human genome, the Telomere-to-Telomere assembly, contains newly identified segmental duplications making up a total of 7% of the total sequence. Using BISER^18^, we re-annotated SDs within the T2T genome, including the Y chromosome **(Supplemental Table 1)**. The sizes of segmental duplications called were between 1kbp and 28Mbp, with 1kbp being the minimum threshold used **(Figure 1C)**. We found that 24.9% of BISER-identified SDs were in regions non-syntenic to the hg38 assembly^19^, suggesting that the T2T assembly is essential to studying these undercharacterized regions.

### Transcription factor binding sites within segmental duplications have high turnover despite sequence similarity

Segmental duplications often copy gene bodies along with their associated *cis*-regulatory elements. Because transcription factor (TF) binding is sequence dependent, we hypothesized that duplicated regions would exhibit similar TF binding profiles due to their shared sequence similarity. To test this hypothesis, we quantified TF binding conservation across segmental duplications. To examine conservation of TF binding sites following segmental duplications, we used Allo to allocate multi-mapped reads in 134 transcription factor ChIP-seq datasets in GM12878, a human B lymphoblastoid cell line **(Supplemental Table 2)**. The GM12878 cell line was chosen as it has a normally presenting karyotype and an abundance of publicly available ChIP-seq datasets. We aimed to avoid issues that arise with large structural variations or polyploidy in other cell lines during our analyses. Additionally, before calling peaks we removed reads that are entirely randomly allocated by Allo to avoid any biases that would come from the use of totally ambiguous regions.

Using this set of TF ChIP-seq experiments, we found the average level of binding site conservation across segmental duplications was 16.5% **(Figure 2A)**, with values ranging between 0% to 39.2% **(Figure 2B)**. Overall, TF binding sites within segmental duplications were not highly conserved regardless of the transcription factor tested. We are confident this low conservation reflects true biology rather than a byproduct of our computational methods. Inaccurate allocation of multi-mapped reads would likely overestimate conservation due to somewhat random assignment across nearly identical regions. To confirm that overall low conservation was not due to the inclusion of multi-mapped reads, we also examined the binding conservation of uniquely mappable peaks at SDs and found the same pattern **(Supplemental Figure 1A)**.

**Figure 2:**
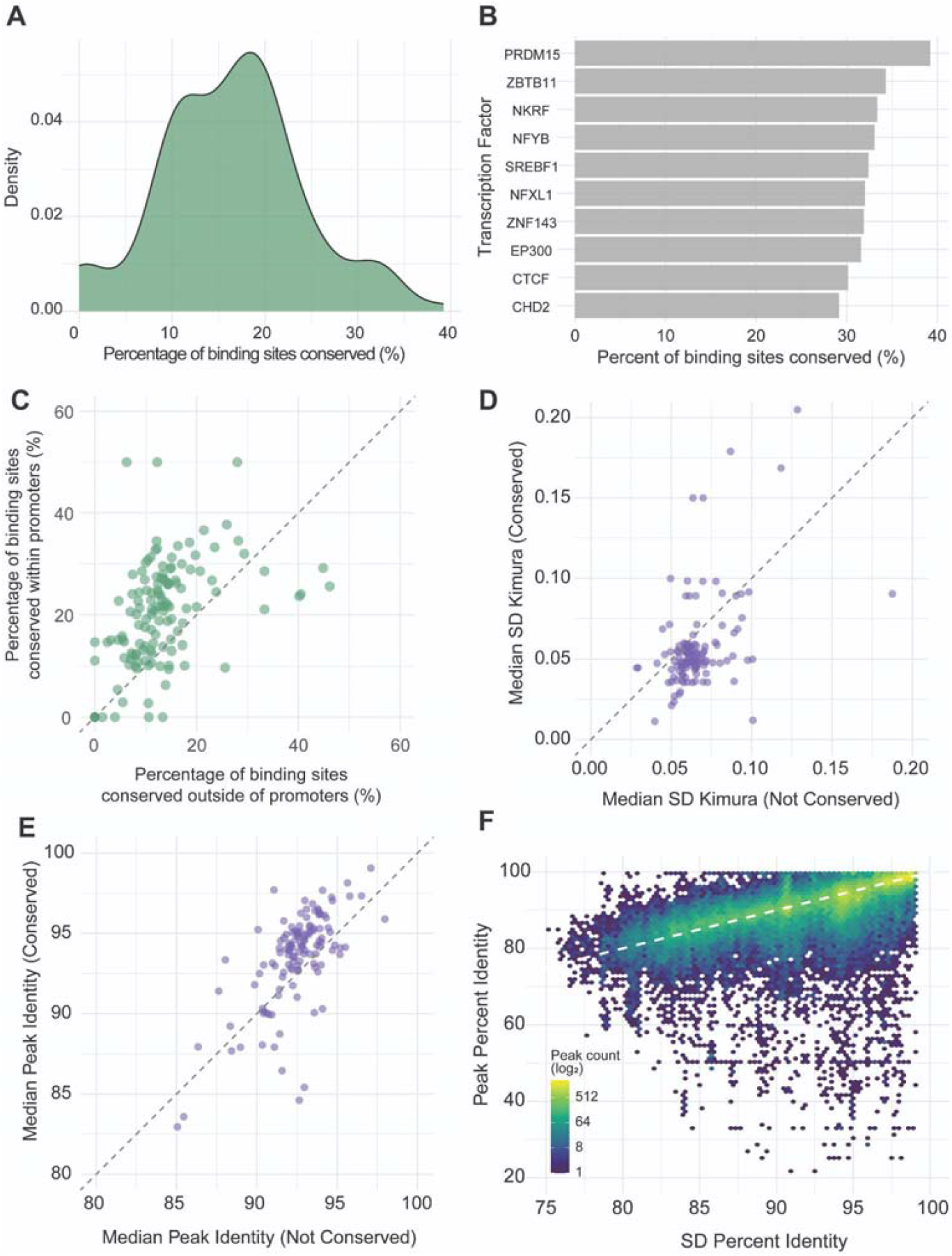
Transcription factor binding sites conservation across segmental duplications in GM12878. **A)** Density plot of the percentage of conservation for binding sites across segmental duplications for 134 transcription factors in GM12878. **B)** Top 10 transcription factors with the highest binding site conservation at segmental duplications. **C)** Comparison between the percentage of binding sites conserved within promoters (0-1kbp upstream of gene start) and percentage of binding sites conserved within distal elements (>1kbp), each point represents one transcription factor. **D)** Comparison between the median Kimura divergence score of associated segmental duplications for conserved and non-conserved peaks for each transcription factor. **E)** Comparison between the median percent identity for conserved and non-conserved peaks for each transcription factor. **F)** Comparison between the percent identity of a peak and the entire associated segmental duplication.

The TF with the highest binding conservation (39.2%) across SDs was PRDM15, which is important for B cell lymphoma metabolism^20^. Other B cell specific factors were also high in conservation relative to other TFs, including PAX5^21^ (24.7% conservation) and ELF1^22^ (24.7% conservation). However, the majority of highly conserved TFs were broadly expressed regulators such as SREBF1^23^ and ZNF143^24^. Other top TFs in our analysis are those ubiquitously expressed and involved in chromatin organization and remodeling such as CTCF^25^, EP300^26^, and CHD2^27^. It is possible that the presence of chromatin remodelers may help maintain chromatin structure after duplication even with the loss of other TF binding. Additionally, many of the TFs with the highest conservation levels are those that are broadly expressed and bind proximally to genes like NFYB^28^ and NKRF^29^. We found that binding sites within promoter regions were more likely to be conserved than those more distal to genes **(Figure 2C)**, emphasizing the importance of regulatory conservation at promoters after duplication. Overall, the most highly conserved TF set was enriched for broadly expressed regulators rather than lineage-specific factors. This finding is consistent with previous studies showing that duplicated genes often evolve tissue-specific expression patterns over time and thus tissue-specific TFs likely have diverging binding patterns after duplication of a binding site.

Next, we investigated how the age of SDs impacted the conservation of transcription factor binding sites. As expected, TF binding sites on older SDs tended to have lower binding site conservation **(Figure 2D)**. As TFs bind in a sequence dependent manner, we also investigated how the percent identity between segmental duplication within TF peaks impacted binding conservation. We found that peaks with higher sequence identity across segmental duplications were more likely to be conserved **(Figure 2E)**, though the difference was minimal. Finally, we hypothesized that TF binding sites would have less divergence than the entire segmental duplication on a sequence level due to the evolutionary pressure to maintain transcriptional control. Surprisingly, we found that the TF binding sites were not more likely to be conserved compared with the entire segmental duplication **(Figure 2F)**. This suggests that the sequence underlying peaks changes at a similar rate to the duplication as a whole. Overall, our findings suggest that TF binding sites are not conserved more than expected on a sequence level, partially explaining their high turnover after duplication.

### Open chromatin is maintained after duplication, while other chromatin states diverge

Histone modifications and other chromatin features both influence and are influenced by TF binding patterns and may therefore be correlated with the high turnover observed at binding sites. Building on our sequence-based analysis of TF binding, we next examined the conservation of chromatin states across segmental duplications. In order to determine chromatin states at SDs, we used DNase-seq and 6 histone modification ChIP-seq datasets to call chromatin states using ChromHMM in 23 cell types **(Supplemental Figure 1B)**. We chose 9 states and named them according to the chromatin marks present at that state as well as their overlaps with genomic annotations **(Figure 3A-B)**.

**Figure 3:**
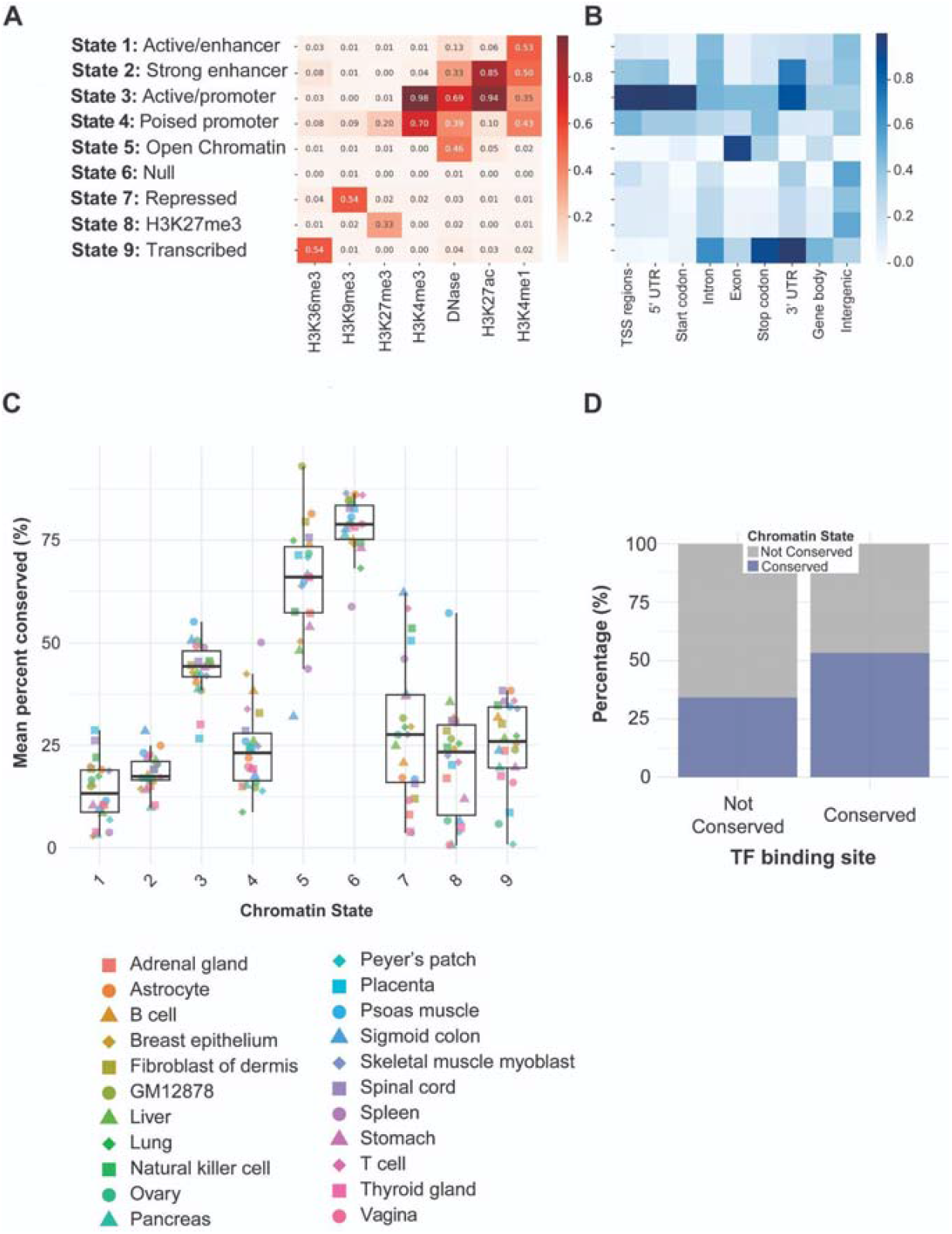
Chromatin states in 23 cell types and their conservation across segmental duplications. **A)** Emission probabilities for each of the 9 chromatin state calls. **B)** Fraction overlap between chromatin state calls in GM12878 and annotated genomic regions. **C)** The mean percentage of 200bp bins that were conserved across segmental duplications for each chromatin state in each cell type. Each data point is the average of that state in one of 23 cell types. **D)** Comparison between the conservation of TF binding sites and the conservation of chromatin states within GM12878.

In examining the retention of chromatin states, we found that, similar to TF binding, a majority of the states were not conserved in any cell type tested **(Figure 3C)**. We found that the states with the highest conservation were the “open chromatin” state 5 and the “quiescent” state 6. In the case of state 6, it is the state that is devoid of any measured accessibility or histone modification signal and makes up a majority of the genome. Therefore, the quiescent state would be expected to be retained by chance across duplicated regions. The relative conservation of the “open chromatin” state (state 5) may be because these areas of the genome are constitutively active areas and thus may contain features like housekeeping genes which are always active. We also saw that promoter-associated state 3 was relatively well conserved. This coincides with our findings that TF binding is more conserved in locations proximal to genes **(Figure 2C)**. We did not see conservation at “weak enhancer” state 1 or “strong enhancer” state 2 suggesting higher turnover at distal elements, coinciding with our results for TFs **(Figure 2C)**. Overall, chromatin states were not maintained after a segmental duplication. The conservation percentages of various chromatin states were similar to that of TF binding sites.

Although TF peaks are generally not conserved **(Figure 2A)**, when a TF binding peak is conserved, the chromatin states are conserved at higher rates **(Figure 3D, Supplemental Figure 2)**. Conversely, sites where the reference locus is bound, but the SD mate is not bound show substantial chromatin state divergence. Most TF-bound peaks are associated with states 2 (“Strong enhancer”), 3 (“Active/promoter”), and 4 (“Poised promoter”) and when the SD mate is not bound, the states transition to a higher proportion of “quiescent” state 6, with few transitions in the opposite direction **(Supplemental Figure 3)**.

Interestingly, the pattern of chromatin state conservation appeared to be dependent on the cell type being analyzed. For example, GM12878 had much higher conservation of open chromatin (state 5) than a majority of the other cell types examined **(Figure 3C)**. It has been previously found that SD regions have higher than expected contributions to the expansion of regulatory elements in immune cell types^30^, possibly explaining why many of the open regions in GM12878 remain open. Additionally, psoas muscle cells had a much higher than average conservation of the H3K27me3-associated state 8, which is characteristic of polycomb-repressed areas of the genome.

### Human-specific segmental duplications are enriched in tissue-specific TF binding sites and show higher turnover than other segmental duplications

As duplicated genes have been shown to contribute to speciation and adaptation in humans^31^, we next compared TF binding sites within human-specific SDs versus those found in other primates. Human-specific SDs were identified using whole-genome alignments of telomere-to-telomere assemblies from six ape species^32^. These species included chimpanzee, bonobo, gorilla, Bornean orangutan, Sumatran orangutan, and siamang. Duplications were classified as human-specific if the corresponding sequence was present exclusively in the human T2T genome and absent from all non-human ape T2T assemblies.

Overall, we found that TF binding sites within human-specific duplications were much less likely to be conserved than those in ancestral duplications **(Figure 4A)**. Similarly, we found that the sequence conservation underlying the TF peaks was lower in human-specific duplications **(Figure 4B)**. It should be noted that ancestral duplications detectable in our analysis may be biased toward those that retained sufficient sequence similarity for alignment and annotation. Rapidly diverging ancestral duplications would be challenging to identify across species and thus are not shown in this analysis. Nevertheless, this pattern suggests that human-specific duplications harbor TF binding sites undergoing rapid regulatory evolution, likely reflecting relaxed selective constraints on duplicated sequences that enable regulatory evolution.

**Figure 4:**
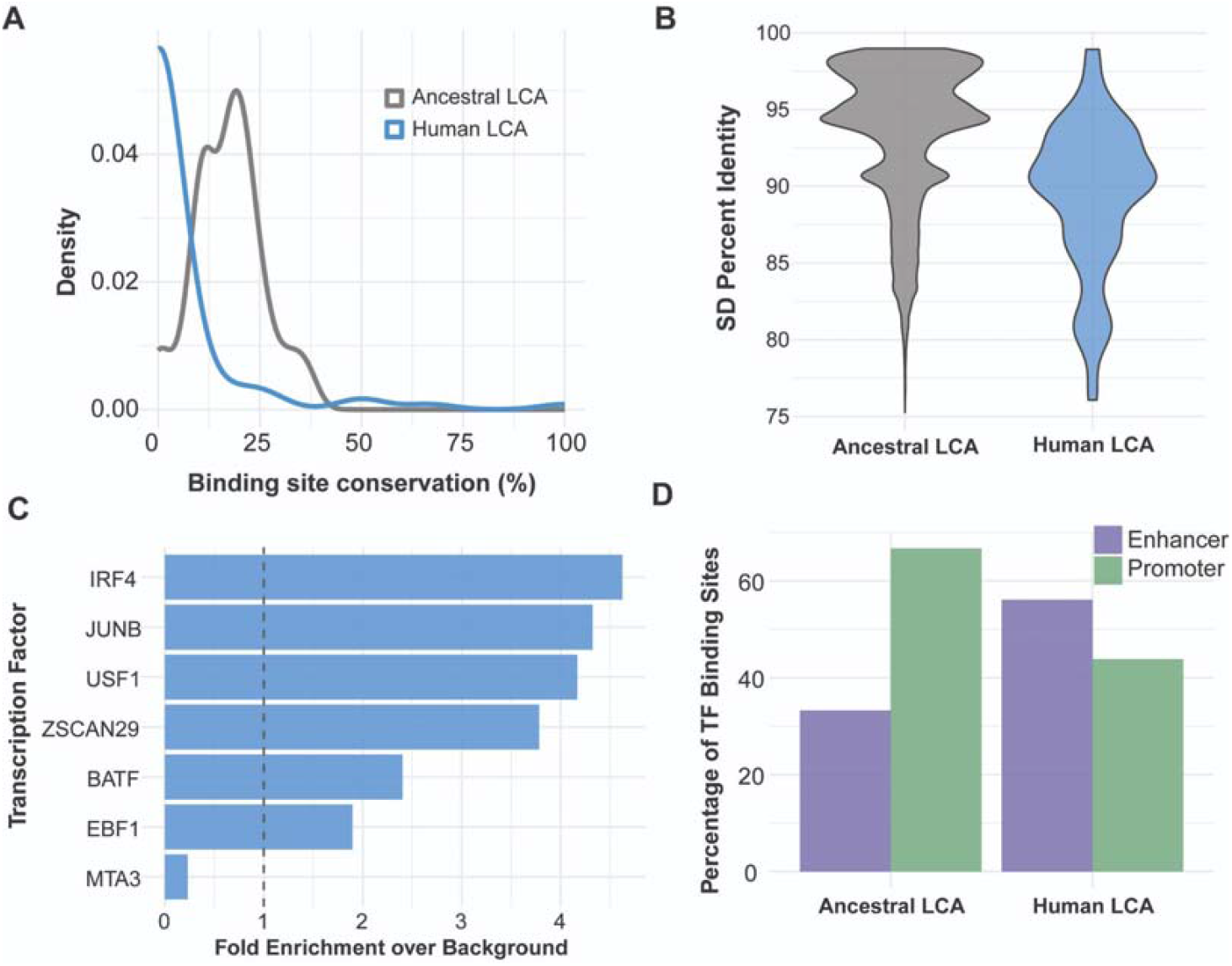
Human-specific segmental duplications display low conservation of their *cis* regulatory regions. **A)** Conservation of TF binding sites across SDs that are human-specific (blue) or ancestral (grey). **B)** The percent identity of SDs in which humans were the last common ancestor (blue) or older primates (grey). **C)** TFs that had the highest enrichment of binding sites on human-specific SDs versus ancestral duplications (background). **D)** The percentage of transcription factor binding sites in GM12878 within enhancers and promoters delineated by the last common ancestor of the SD the binding site was on.

As previously discussed, it has been found that SD regions have higher than expected contributions to the expansion of regulatory elements in immune cell types^30^. We analyzed which TFs in GM12878 had the highest concentration of binding sites within human-specific SDs **(Figure 4C)**. We found that TFs involved in immune cell specific regulation, such as IRF4, JUNB, BATF, and EBF1, were enriched within these human-specific SDs. Our results suggest that human-specific duplications are enriched in lineage specific factors whose binding quickly diverges.

We next analyzed whether human-specific duplications were enriched with specific *cis* regulatory regions, such as enhancers or promoters **(Figure 4D)**. We found that ancestral duplications tended to have a higher representation of promoter-associated states 3 and 4 and human-specific duplications tended to have a higher representation of enhancer-associated states 1 and 2. This enrichment of enhancers in human-specific duplications is consistent with their reduced conservation, as enhancers typically exhibit greater evolutionary flexibility than promoters and serve as hotspots for regulatory diversification.

### Despite regulatory turnover, segmentally duplicated genes have similar expression patterns

Previous studies have shown that paralogous genes exhibit correlated expression compared with randomly selected gene sets^33^. Focusing specifically on SD paralogs, we used RNA-seq data from 23 cell types to assess expression conservation. Consistent with prior work, paralogs overall showed significantly higher average expression correlation than either random gene pairs (Wilcoxon rank-sum adj p = 5.63×10□^91^) or genes drawn from the same gene ontology categories (Wilcoxon rank-sum adj p = 4.01×10□^69^) **(Figure 5A)**. Notably, SD paralogs exhibited even higher expression correlation across tissues than the full set of paralogs (Wilcoxon rank-sum adj p = 9.12×10□^65^). This observation is expected, as SD paralogs represent the youngest paralogous pairs. In addition, higher sequence similarity within their gene bodies suggests reduced functional specialization relative to older paralogs, for which specialization often leads to tissue-specific expression patterns^34^.

**Figure 5:**
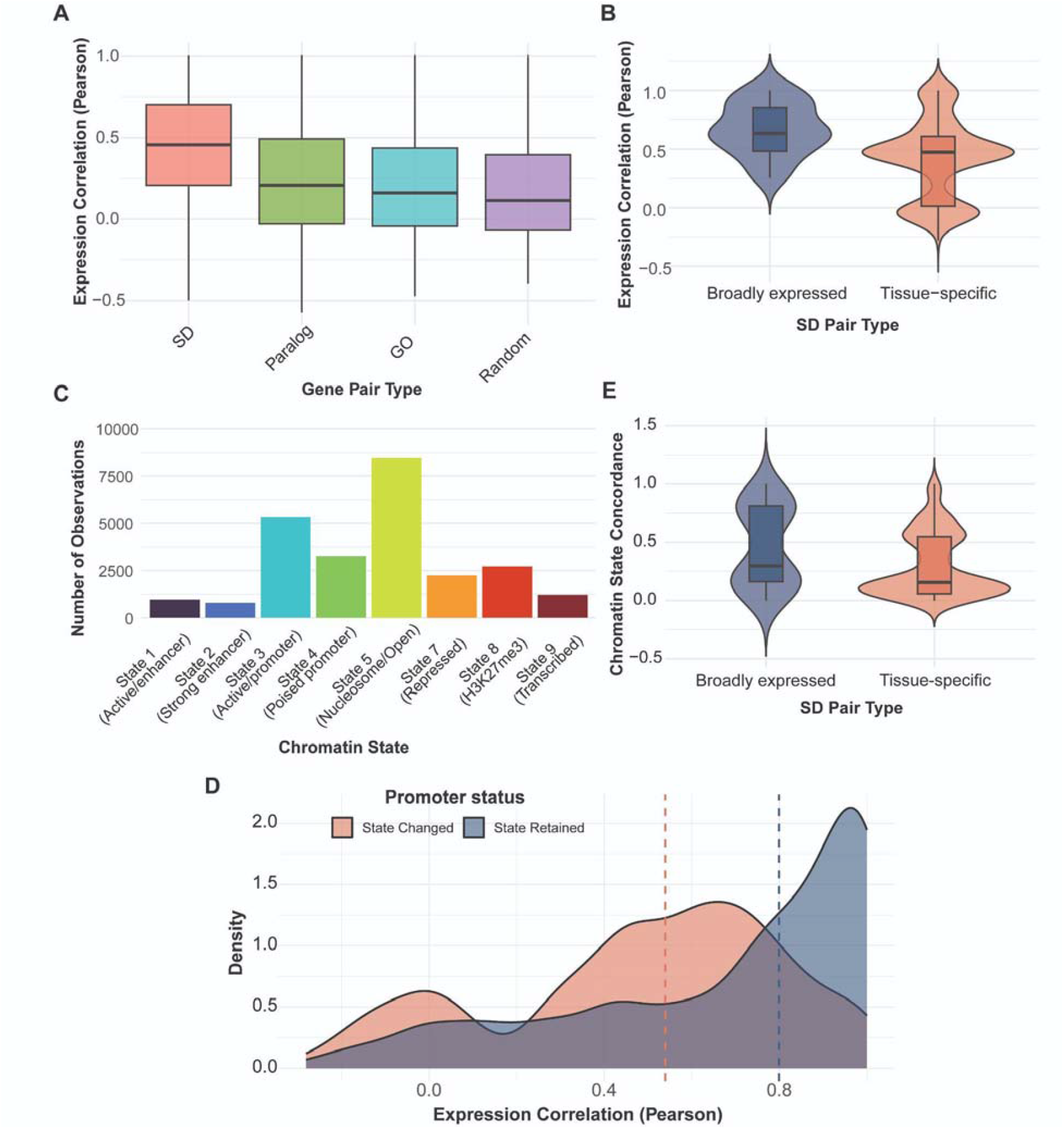
Gene expression is conserved between segmentally duplicated genes across 23 cell types. **A)** Pearson correlation coefficients of gene expression for gene pairs in categories including random pairs, pairs of genes with the same gene ontology, genes in the same paralogous family, paralogous gene pairs from segmental duplications. **B)** Pearson correlation coefficient of the gene expression of segmentally duplicated gene pairs in which both genes are broadly expressed (15/23 cell types) or tissue-specific (3 or less cell types). **C)** Number of promoters with each chromatin state across 23 cell types, each promoter is represented 23 times. **D)** Chromatin state concordance between the promoters of segmentally duplicated gene pairs in which both genes are broadly expressed (15/23 cell types) or tissue-specific (3 or less cell types). **E)** Density plot showing the Pearson correlation coefficient of gene expression vectors for gene pairs with the same chromatin state in their promoter (blue) or different states at their promoter (red).

To assess the impact of tissue specificity on SD paralog expression, we classified genes as tissue-specific (expressed in three or fewer cell types) or broadly expressed (expressed in at least 15 cell types). Tissue-specific SD paralogs showed greater expression divergence than broadly expressed SD paralogs **(Figure 5B)**. This reduced correlation parallels our previous findings that tissue-specific transcription factors exhibit higher regulatory turnover than broadly expressed TFs.

We next examined chromatin states at SD paralog promoters across cell types **(Figure 5C)** (Wilcoxon rank-sum adj p = 7.9×10□^6^). As expected, promoter-associated states (states 3 and 4) and the open chromatin state (state 5) were the most prevalent. We then tested whether promoter chromatin state concordance was associated with expression similarity. SD paralogs whose promoters shared the same chromatin state generally displayed higher expression correlation **(Figure 5D)** (Wilcoxon rank-sum adj p = 4.58×10^-82^). Given the influence of tissue specificity on expression correlation, we hypothesized that tissue-specific genes would also show reduced chromatin state concordance across cell types **(Figure 5E)**. Although this difference was significant (Wilcoxon rank-sum adj p = 4.38×10□^3^), its magnitude was smaller than that observed for expression correlation, likely reflecting the more limited quantitative resolution of chromatin state annotations for predicting gene expression.

### Spatial proximity may buffer transcriptional divergence when TF binding sites are lost during duplication

While we have seen that TF binding and chromatin states are not conserved at high rates between segmentally duplicated regions of the genes, we also confirmed that their expression correlation was still higher than expected. Previous studies have shown that paralogous genes (including those not from SDs) are closer within the three-dimensional genome than expected and may share distal regulatory elements^11,35^. We therefore hypothesized that SD regions of the genome form shared regulatory networks through close contact with the same distal elements as their ancestral duplication.

To examine the contact intensity of SD paralogs, we analyzed Hi-C data from 15 of the above cell types. In this analysis, we limited it to only uniquely-mappable SD paralogs to get the highest confidence contacts and estimated the contact frequencies using Cooler^36^ **(Supplemental Figure 4A)**. As many SD regions can be tandemly duplicated and thus would automatically show high contact intensity, our random background set was constructed to have random region pairs of the same size and linear distance as the paralog in question **(Supplemental Figure 4B)**. We also validated our pipeline showing that closer pairs had higher contact intensity **(Supplemental Figure 4C)**.

As anticipated, we found that the contact intensity of SD paralogs was significantly higher than random distance-controlled regions in all cell types analyzed **(Figure 6A)**. Additionally, we found that low TF conservation pairs tended to have the highest contact among all pairs **(Figure 6A)**, though low TF conservation pairs also tended to be linearly closer than high TF conservation pairs **(Figure 6B)**. In comparing the Hi-C contact intensity between SD gene pairs to the correlation of their expression, all interactions were significant regardless of TF binding site conservation **(Figure 6C)**. Together, these results suggest that enhanced spatial proximity between SD genes may partially buffer transcriptional divergence following the loss of shared TF binding by facilitating access to common distal regulatory elements.

**Figure 6:**
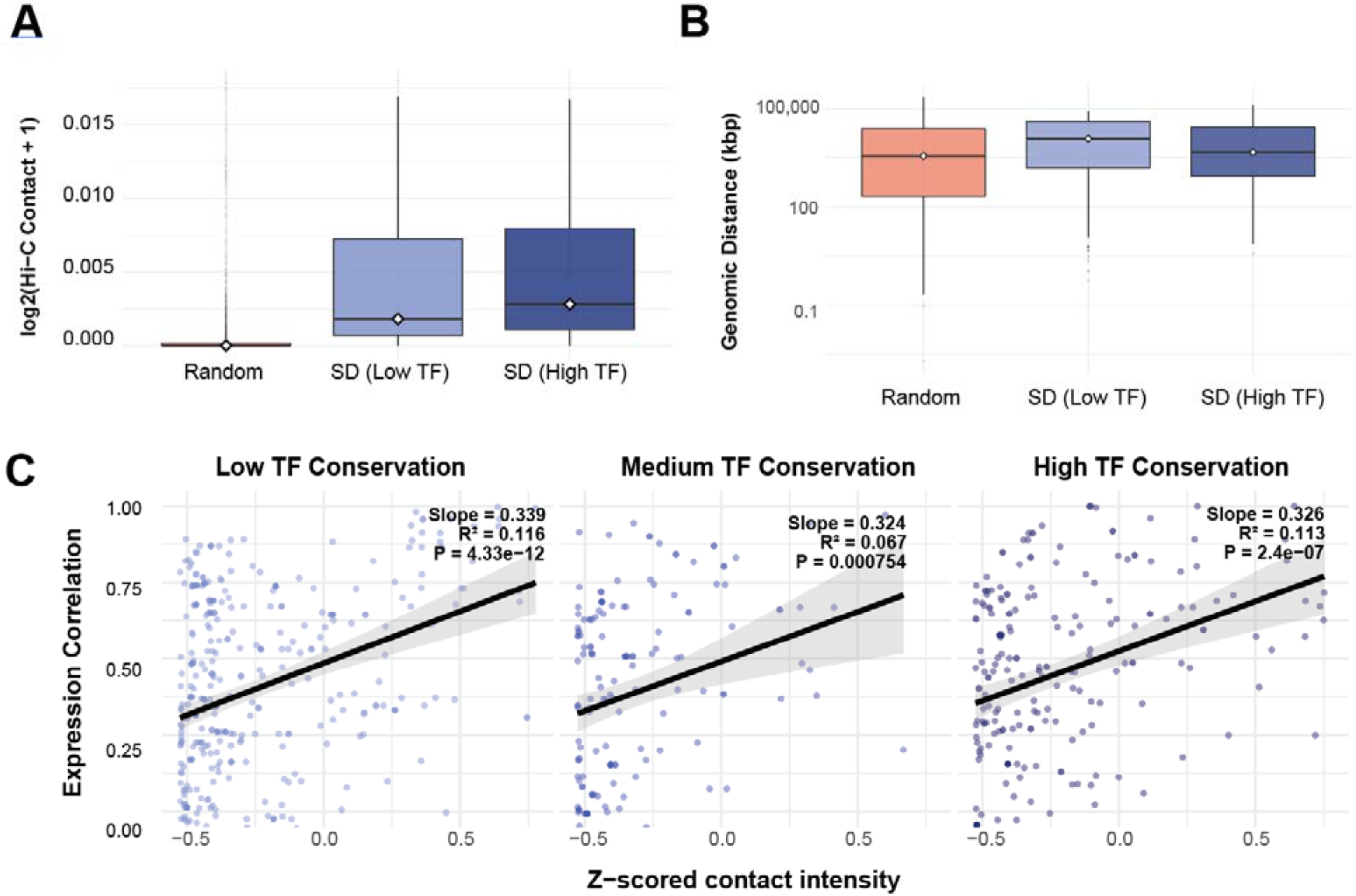
Contact frequencies of gene pairs within segmental duplications based transcription factor binding. **A)** Log_2_ contact frequencies of random gene pairs versus segmentally duplicated regions with high (≥75th percentile) or low TF conservation (≤25th percentile). **B)** Genomic distances between random gene pairs versus segmentally duplicated regions with high (≥75th percentile) or low TF conservation (≤25th percentile). **C)** Relationship between chromatin contact frequency and expression correlation for segmentally duplicated gene pairs, separated by transcription factor binding site conservation. Gene pairs are categorized by TF conservation: low (≤25th percentile), medium (25th-75th percentile), and high (≥75th percentile). Figure excludes the top 10% of outliers for visualization.

## Discussion

The evolution of genome complexity is fundamentally driven by gene duplication, yet the regulatory mechanisms governing these duplications have remained understudied due to the limitations of short-read sequencing assays and incomplete reference genomes. By leveraging the Telomere-to-Telomere human genome assembly and probabilistically allocating multi-mapped reads, this study provides an overview of the regulatory landscape of segmentally duplicated regions within the human genome. Our findings revealed an unexpected result. While SD regions maintain high sequence identity, their regulatory profiles show rapid divergence. Additionally, in many cases, the expression of SD paralogs was partially maintained even with the loss of regulatory profiles.

On an evolutionary scale, our results show that the sequences underlying TF peaks evolve at approximately the same rate as the rest of the SD **(Figure 2F)**. This lack of purifying selection at most duplicated binding sites implies an immediate period of relaxed evolutionary constraint following duplication. The relaxed constraints on these regions may allow for rapid regulatory subfunctionalization and neofunctionalization, where the two copies of a gene begin to be regulated by different sets of TFs. The divergence in regulation potentially sets the stage for tissue-specific specialization and expression. Our analysis of human-specific SDs reinforces the role of duplication in recent evolution. These regions showed even lower TF binding conservation and a distinct enrichment for lineage-specific TFs (such as IRF4 and EBF1) as well as an enrichment for distal regulatory elements such as enhancers.

While TF binding and chromatin states displayed high turnover almost immediately after duplication, it appears that gene expression patterns are still partially maintained. One plausible explanation for the high correlation between these paralogs was likely their three-dimensional proximity. The spatial proximity of SDs allow them to share a regulatory neighborhood, possibly allowing access to the same distal enhancers. Even if one copy loses a specific binding site, its physical proximity to the other copy and its associated regulatory elements may maintain its transcriptional activity. Such spatial buffering would provide a safety net, allowing for the observed regulatory turnover without a detrimental loss of gene expression. Spatial buffering may therefore facilitate the eventual neofunctionalization or subfunctionalization of duplicated genes.

In summary, we show that the regulatory environment of SDs is highly dynamic. The transition from a duplicated sequence to a functionally distinct gene involves a rapid loss of ancestral regulatory elements that is possibly balanced by the sharing of distal regulatory elements. Future studies using long-read technology with spatial genomics may better elucidate the interactions between segmentally duplicated regions of the genome.

## Methods

### Datasets and code availability

Datasets used in this study were downloaded from ENCODE^37^. Identifiers for the chromatin state analysis **(Supplemental Table 3 and 4)**, RNA-seq analysis **(Supplemental Table 5)**, GM12878 ChIP-seq analysis **(Supplemental Table 2)**, and Hi-C analysis **(Supplemental Table 6)** can all be found in supplemental materials. All available code for this project can be found in the following GitHub repository: https://github.com/seqcode/segdups_project

### Overlaps with annotations

To find the overlap with repetitive elements, the annotations from RepeatMasker^38^ for the T2T v2.0 (January 2022) genome were downloaded from the UCSC genome browser. Syntenic and non-syntenic region annotations were taken from Aganezov et al. 2022^19^. Annotations for segmental duplications are detailed in the below section. Overlaps between reads and each annotation were performed using BEDTools^39^ v2.31.1 intersect (-u).

### Segmental duplication identification and evolutionary distance calculations

BISER^18^ v1.4 was used to identify segmentally duplicated regions in the T2T genome v2.0 (softmasked) using default settings. The resulting file was filtered by segmental duplications that had less than a 25% error rate or 75% identity (column 8 of the BISER output). Additionally, the BED file used as the annotation for SDs was constructed so the alignment went in both directions. The CIGAR string output by BISER was used to construct alignments for all same-stranded segmental duplications along with BEDTools v2.31.1 getFasta and the T2T genome. A custom python script was used to create a dictionary that contained information about predicted segmental duplications including their percent identity and Kimura 2-parameter^40^ score as well as the nucleotide content at each paired duplication. The script can be found in the GitHub repository under “create_dict.py”. A compressed version of the Python dictionary can be retrieved from figshare at the following location: “https://doi.org/10.6084/m9.figshare.28458767.v1“.

### DNase-seq and ChIP-seq pre-processing

To more accurately analyze the data, we used different aligners based on the types of reads they were optimized to handle. Bowtie is more accurate for shorter reads whereas Bowtie2 is most accurate for paired-end reads. For this reason, single-end reads under 50bp were aligned to the T2T genome (chm13 v2.0) using Bowtie^41^ v1.0.0 with the arguments “--best --strata -m 25 -k 25 --chunkmbs 1024”. Single-end reads equal to or over 50bp were aligned to the T2T genome using Bowtie2 v2.5.0 with the argument “-k 25”. For paired-end reads, regardless of length, Bowtie2 v2.5.0 was used for alignment with the arguments “-k 25 --no-mixed --no-discordant”. Allo was used to allocate multi-mapped reads using default settings with the exception of using the “--mixed” argument for all histone ChIP-seq datasets analyzed. Reads allocated by Allo with the ZZ tag, indicating random allocation, were filtered from our analyses to avoid issues with fully identical regions. Following this, peaks were called on ChIP-seq replicates separately using MACS2^42^ v3.3.0 with the argument “-p 1e-3” to raise to p-value threshold and “--broad” for all histone modification datasets analyzed. To combine replicates, irreproducible discovery rate (IDR)^43^ was used with default settings.

### ChIP-seq conservation analysis

To investigate the conservation of ChIP-seq peaks, a custom Python script was created. Briefly, the peaks called in 134 GM12878 ChIP-seq datasets as well as the dictionary detailed above were used to assign peaks to SDs. If a segment was duplicated more than once, each pair was treated as a separate measurement. Additionally, the script calculated the percent identity under the peak as well as the Kimura 2-parameter score. These values, along with those for the entire SD were in the final output. The Python script can be found in the GitHub repo under the name “peak_comps.py.”

### Comparative genomics analysis

A whole genome alignment of size ape Telomere-to-Telomere genome assemblies was downloaded in the form of a HAL file^32^. These species included chimpanzee, bonobo, gorilla, Bornean orangutan, Sumatran orangutan, and siamang. A segmental duplication was considered human-specific if it was only present in the human genome. Otherwise, the segmental duplication was considered ancestral.

### Chromatin state calling

We used ChromHMM^44^ v1.23 to perform chromatin state analysis across all 23 cell types using the same histone modification ChIP-seq datasets. ChromHMM was run iteratively with state numbers ranging from 8 to 15. The resulting models were compared using the CompareModels functionality in ChromHMM to see how well models with various numbers of chromatin states correlate with each other. The optimal number of chromatin states was determined by selecting the model that minimized redundancy while maximizing the identification of unique chromatin states. Each chromatin state was named to reflect possible biological activity based on the emissions profile for that state.

To analyze the conservation of each chromatin state, a similar method was deployed to the one used for the TF peak sets. First, the 9 state ChromHMM output was split into 200bp bins for the entire genome. Following this, the conservation of each bin was determined exactly as it was for the ChIP-seq datasets using the “peak_comps.py” script. For plotting the conservation of each state per cell type, the average was calculated across all 200bp bins in the genome with each specific state call.

### RNA-seq analysis

Reads were aligned to the T2T genome (chm13 v2.0) using STAR^45^ v2.5.2 with the arguments “--outSAMtype BAM Unsorted --outSAMmultNmax 25 --outFilterType BySJout” to allow for the retention of multi-mapped reads. Following this, Allo v1.2 was used to allocate the reads using the arguments “--readcount --splice”. These settings in Allo allocate the multi-mapped reads solely based on the uniquely mapped reads in a 500bp region with introns removed. Following this, featureCounts^46^ v2.0.2 was used with the arguments “-M -t exon -g gene_id”. For each software listed in the pipeline, paired-end settings were utilized for paired-end datasets. Following the use of featureCounts for each cell type, the counts tables for all samples were merged for the final analysis.

Using the summarized counts table, genes were removed from the analysis that did not have at least 50 counts summed across all samples. The mean between biological replicates of each cell type was used for the remaining analyses. The table was then merged with a paralog annotation file obtained with permission from GeneCards^47^. In order to identify the segmentally duplicated paralogs, a BED file was downloaded with gene locations within T2T from the UCSC genome browser^48^ (retrieved Jan 2025).

Next, the dictionary above was used to identify genes within segmentally duplicated regions and their possible paralogs. Gene B was identified as a paralog of Gene A if Gene B covered at least 80% of Gene A in an identified segmental duplication. To save time, only every 1kb was checked for matches for each gene as opposed to the entire gene. The script for this can be found under the name “seg_pairs.py” in the GitHub repo for this project. The pairs identified via this analysis are in **Supplemental Table 7**. Pairwise comparisons were done using Wilcoxon rank-sum tests with Benjamini-Hochberg correction.

### Hi-C analysis

Pre-processed pairs files were downloaded from ENCODE for 15 different cell types **(Supplemental Table 6)**. The pairs files were lifted from hg38 to T2T using HiCLift^49^ v1.0. The chain file was downloaded from the T2T consortium’s annotations (retrieved Dec 2025)^50^. A custom script utilizing Cooler^36^ was used to calculate the contact frequencies between regions of the genome. Briefly, for each pair of genomic regions, it generates a corresponding random pair that maintains the same genomic distance. The script then calculates the mean contact frequency between each pair of regions across all cell types using mcool files at 10kb resolution, processing both actual and random pairs.

## Acknowledgements

This work was supported by the National Institutes of Health grant R35GM144135 (to S.M.) and the National Science Foundation DBI CAREER 2045500 (to S.M.). A.M. is currently supported by the National Science Foundation (NSF) through the NSF National Synthesis Center for Emergence in the Molecular and Cellular Sciences (NCEMS) under the National Science Foundation grant MCB-2335029. Any opinions, findings, and conclusions or recommendations expressed in this material are those of the authors and do not necessarily reflect the views of the National Science Foundation or NCEMS. A. B. was partially supported by the National Institutes of Health training grant T32 GM152354 and by a GlaxoSmithKline Graduate Fellowship. J.Y. was partially supported by Rising Researcher award ICDS_RR25_027636 from Penn State’s Institute for Computational & Data Sciences (RRID:SCR_025154). The authors thank the members of the Center for Eukaryotic Gene Regulation at Penn State for helpful feedback and discussions.

## Notes

### Competing Interest Statement

The authors have declared no competing interest.

