## Supplementary Figures for "Widespread Regulatory Turnover Across Human Segmental Duplications"

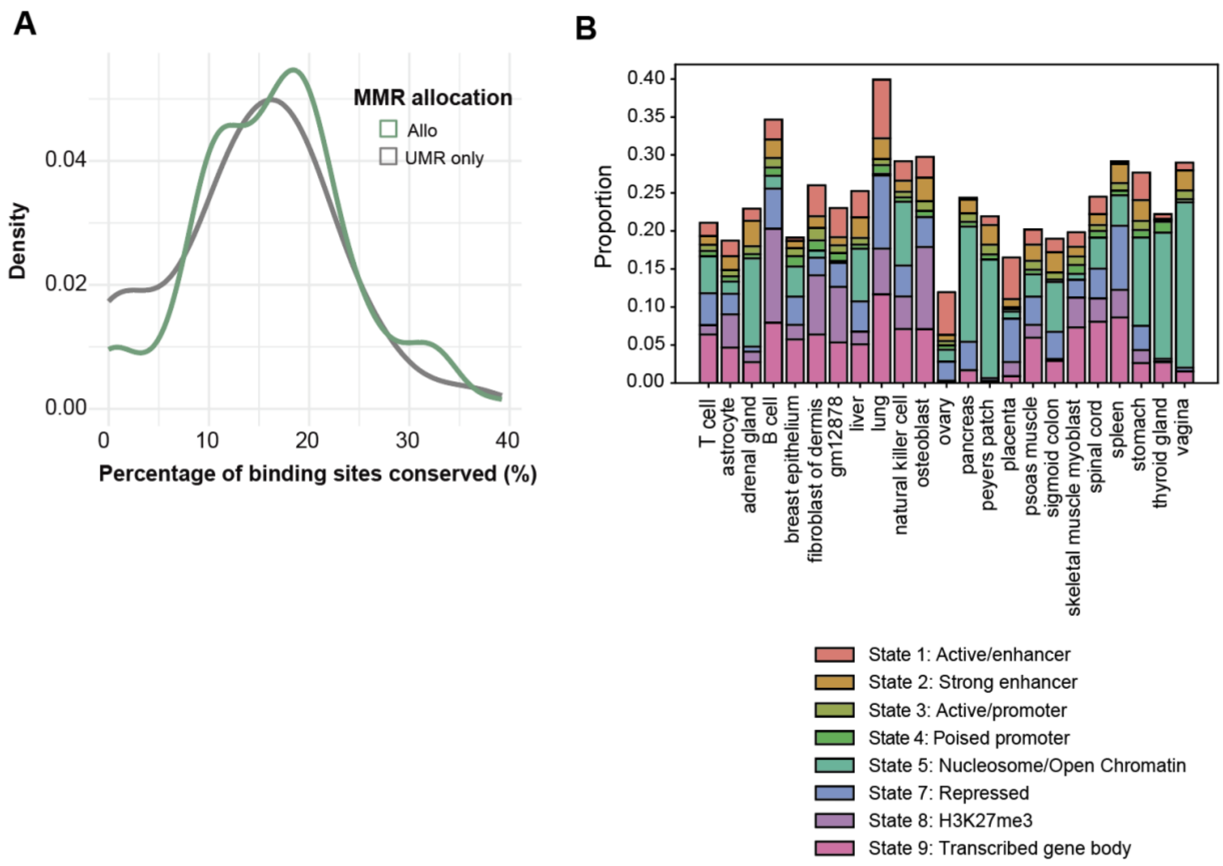


**Supplemental Figure 1:**

**A)** Percentage of binding sites conserved across GM12878 ChIP-seq datasets before and after allocating multi-mapped reads with Allo **B)** Fraction of genome (disregarding null state ) that had each chromatin state in 23 cell types.

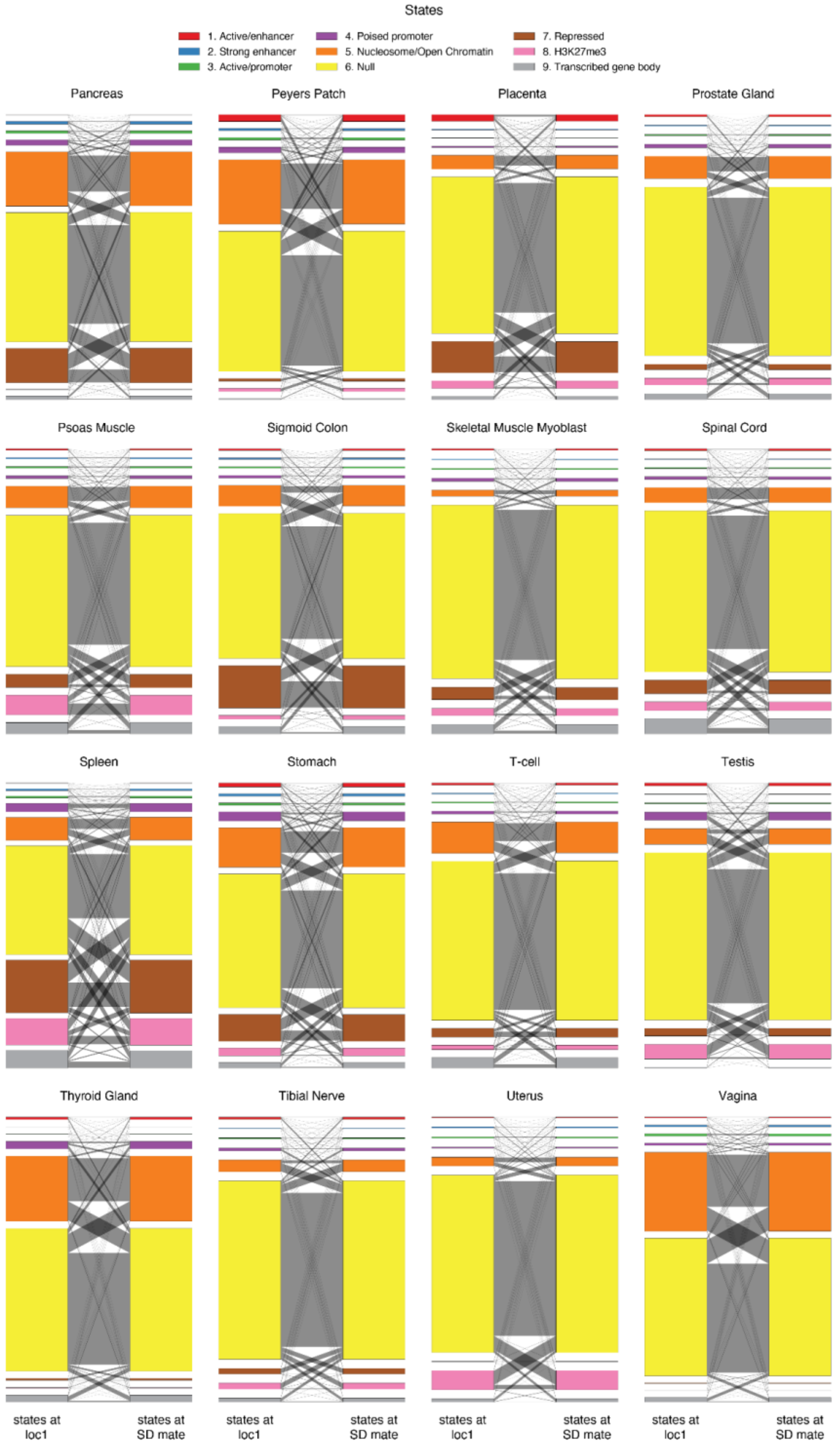


**Supplemental Figure 2:**
Flow plots summarizing chromatin state transitions over segmental duplications in 16 different cell types. Within each category, the height of the colored bar is proportional to the number of peaks annotated with that chromatin state. Gray ribbons connect states between loci with the widths proportional to the number of states transitioning from one state to another.

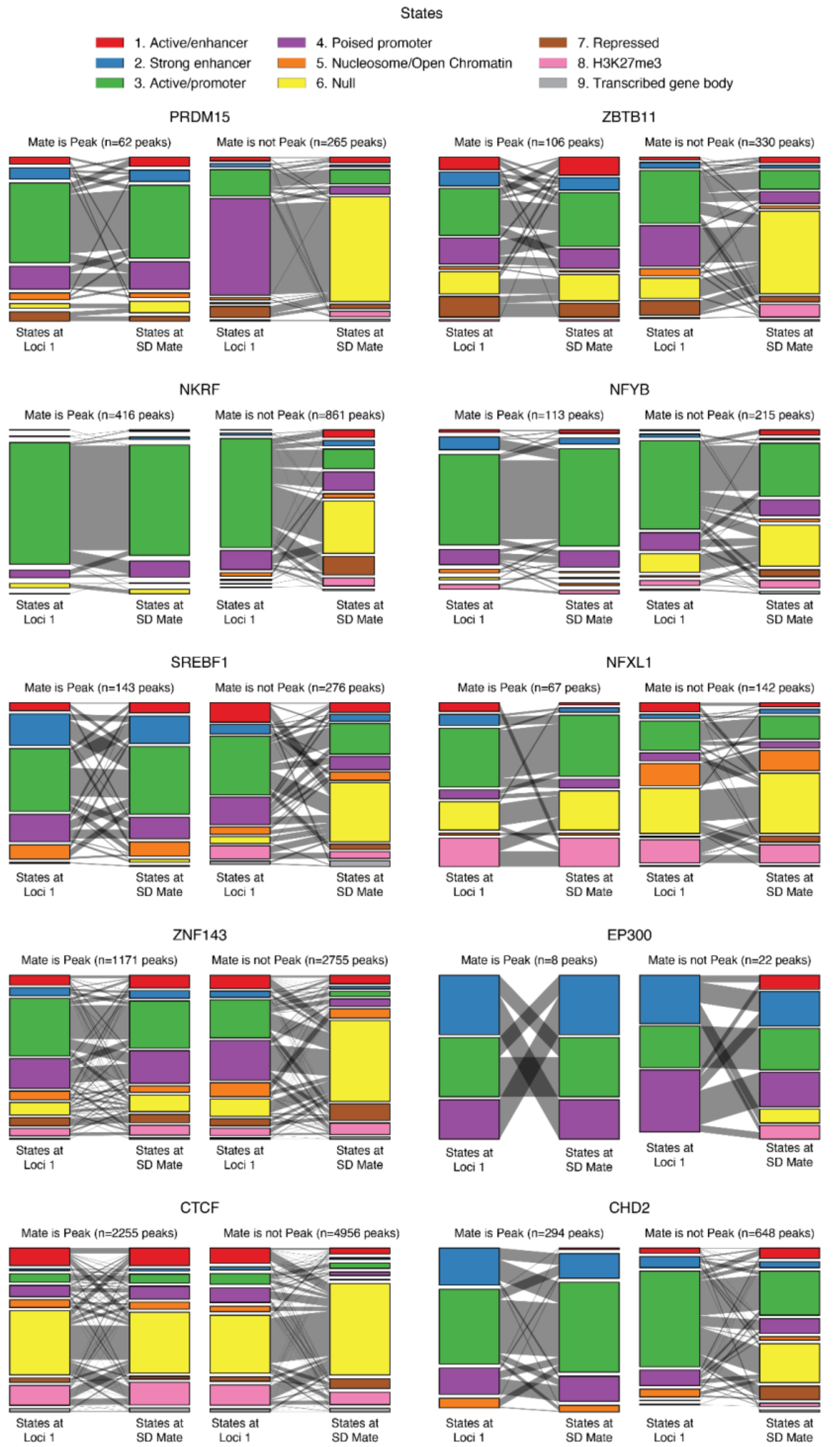


**Supplemental Figure 3:**
Flow plots summarizing chromatin state transitions at peaks for the top 10 most conserved TFs. For each TFs, peaks are stratified into two categories: conserved peaks (left pair; "Mate is Peak") and non-conserved peaks (right pair; "Mate is not Peak"). Within each category, the height of the colored bar is proportional to the number of peaks annotated with that chromatin state at the reference locus ("States at Loci 1") and the SD site ("States at SD Mate"). Gray ribbons connect states between loci with the widths proportional to the number of states transitioning from one state to another. The total number of peaks in each category is indicated above each panel.

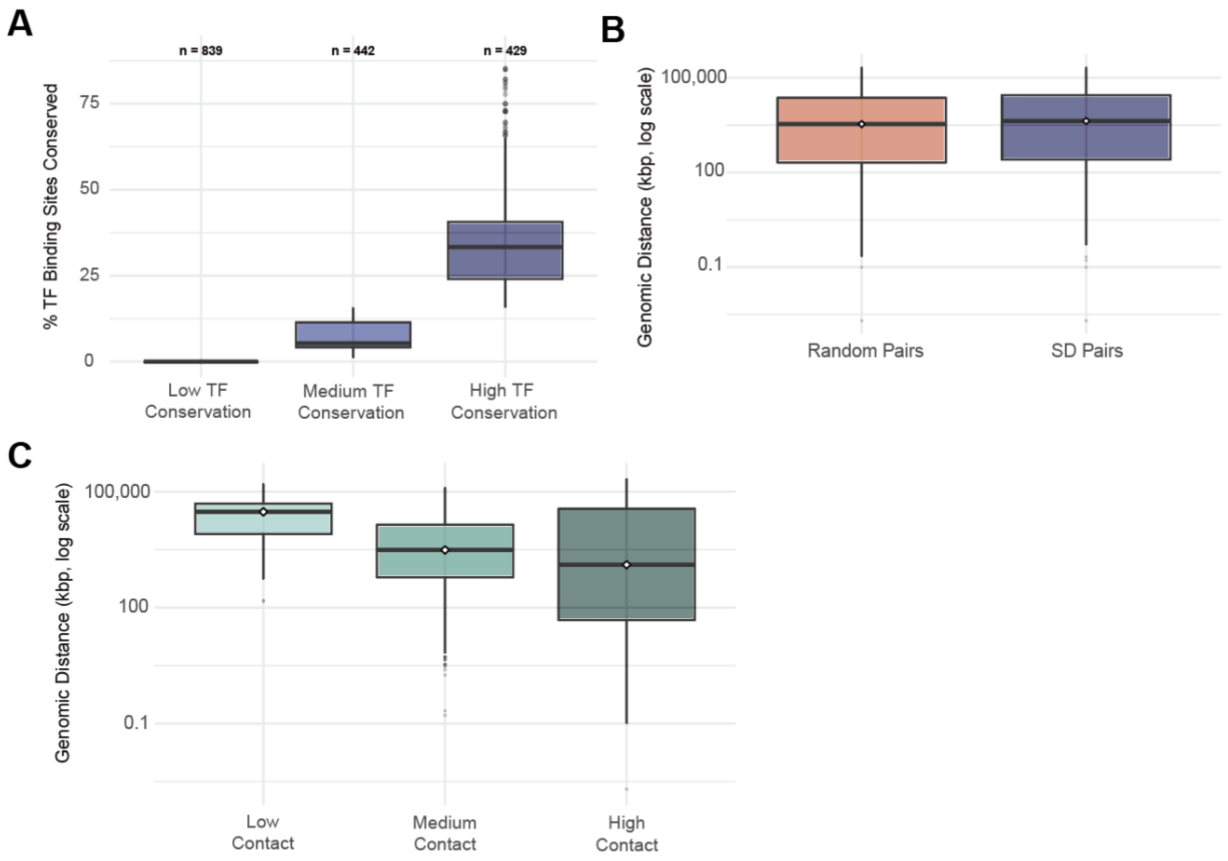


**Supplemental Figure 4**

**A)** Transcription factor binding site conservation for three distinct categories: low (≤25th percentile), medium (25th-75th percentile), and high (≥75th percentile). **B)** Genomic distances between segmentally duplicated gene pairs and random size matched regions. **C)** The relationship between genomic distance in basepairs (log scaled) compared to the calculated contact frequency of region pairs using Cooler.
